# *Corythauma ayyari* (Insecta, Heteroptera, Tingidae) depends on its host plant to invade Europe

**DOI:** 10.1101/2023.11.15.567321

**Authors:** Manon Durand, Eric Guilbert

## Abstract

Biological invasions increase with the intensity of globalization, human activities, and climate change. Insects represent a high potential of invasive species due to their adaptability to new environment. We analysed here the invasiveness of an Asian phytophagous bug, *Corythauma ayyari* (Heteroptera, Tingidae), recently recorded in Europe, and that depends on *Jasminum* spp., an ornamental plant widespread in Europe. We modelled its current distribution, projected it into the future and tested its niche overlap between native and invaded areas. When considering the host plant as environmental variable, the analysis shows that *C. ayyari* shifted to a new ecological niche but its distribution is restricted by its host plant distribution. Including or excluding the host plant as environmental variable has an impact on species distribution and should be considered when dealing with phytophagous species.

## Introduction

An invasive species is an exotic species that begins to proliferate in a new geographical habitat. Competing with native species, the proliferation of this invasive species can lead to disruptions in the functioning of the host ecosystem and can the source of serious ecological, economic and health impacts.

While the number of biological invasions continues to increase, particularly with the intensity of globalization and the acceleration of climate change, they are now one of the main causes of the current loss of biodiversity. Trade would be the major driver of alien species introductions at least for invertebrates and algae [1]. In order to fight against their proliferation and the damages, it is necessary to identify these species and to analyse their invasive potential.

Insects are the most diverse group of living species on earth, thanks in part to their ability to adapt to new environmental conditions. It is therefore not surprising that many insect species are considered invasive or potentially invasive. For example, 50% of recently invasive species belong to the order Hemiptera [2]. *Corythauma ayyari* (Drake) belongs to the family Tingidae (Hemiptera, Heteroptera). It has been first described from India (Drake 1933) and found on *Jasminum pubescens* (Retz.), today synonymized with *J. multiflorum* (Burm.). It is notably found in Pakistan, Sri Lanka, Laos, Thailand, Malaysia, India, Indonesia, and Singapore [2, 3, 4] It is known as a pest of Jasmine in Southern India [5]. Recent observations report its presence in France [4, 6], Italy [3, 4, 7], Spain [2], Tunisia [8], Israel [9], and Syria [10]. Most records mention the three main host plants *Jasminum officinale* L., *J. grandiflorum* L., and *J. sambac* L. (Oleaceae) but also *Volkameria inermis* L. (Lamiaceae) [10] and *Trachelospermum* sp. (Apocynaceae) [6].

It is hypothesized to have been accidentally imported via trade in of its host plants, which are ornamental plants widely used in public and private gardens, as they are highly valued for their fragrance [2]. However, the maintenance of *C. ayyari* on Jasmine species is not without damage to the host plant. It has been observed that adults and nymphs develop on the underside of the leaves and feed on sap. The excreta of the tingid mar the leaves, greatly decreasing photosynthesis, which deteriorates the palisade parenchyma and leads to progressive desiccation of the plant [2–4, 7, 10]. The recent arrival in Europe of *C. ayyari* suggests that it could wreak havoc on Jasmine species but also other ornamental and/or crop plants, causing significant environmental and economic damage.

Species Distribution Models (SDM) are increasingly used to answer biogeographic and ecological questions. Indeed, these models allow to estimate the probability of presence of a species according to different biotic and abiotic variables, in order to expose the favorable areas for its presence [11, 12]. These tools are particularly interesting when studying invasive or potentially invasive species. Previous studies have indeed demonstrated the interest and reliability of these methods to quantify the probability of presence and niche differences of an invasive species [13, 14].

Phytophagous insects such as Tingidae which are sap suckers, are strictly dependent of their host plant. Particularly if they are monophagous. Therefore, their distribution highly depends on the distribution of the host plant. Two publications have already focused on the current distribution of *C. ayyari* at the level of the Mediterranean basin and specially in the Iberian Peninsula [2] and in Italy [4]. Therefore, it makes sense to reinforce them by focusing here on the invasive potential of the species.

The aim of this study is to analyse the invasive potential of *C. ayyari* by modelling its current distribution and its potential ecological niche in Europe, according to the current climatic conditions and the distribution of its main host plants. We modelize *C. ayyari* distribution including or not the four main host plants (*J. officinale*, *J. multiflorum*, *J. grandiflorum* and *J. sambac*), as environmental variable to evaluate their influence on *C. ayyari* distribution. On this basis, *C. ayyari* potential distribution is projected into the future. The native and the invasive ecological niches are compared to see if a shift from native to invasive niche is observed.

## Materials & Methods

### Study site

The study focuses on the potential area of invasion of *C. ayyari*, according to the distribution of its host plants and considering their native area in Asia. It includes an area between 18° West to 141° East and between 1° to 70° North; covering the invaded area in Europe and the Mediterranean Basin between 18° West to 40° East and 30° to 70° North, and the native area which covers a surface within 1° to 46° North and 65° to 141° East. It covers also part of Africa, where occurrences of Jasminum species are known. If occurrences of the host plants are known from North America, none of *C. ayyari* is known there.

### Species occurrences

Occurrences of *C. ayyari* were retrieved from the GBIF network (dio:10.15468/dl.b3jj26) and from scientific publications [4, 6, 8, 15, 16] and also from personal communications. Similarly, occurrences of *J. officinale*, *J. grandiflorum*, *J. multiflorum,* and *J. sambac*, were extracted from the GBIF network (<u>dio:10.15468/dl.9sbc2q</u> and <u>dio:10.15468/dl.2j7zqa</u>). *Jasminum officinale* has an extended distribution in Europe, compared to the three other *Jasminum* species. There are only 21 records of *J. grandiflorum* in South West Europe, the rest are located in Asia. *Jasminum multiflorum* is sparcely recorded from South East Asia and totalizes two records in Europe (Germany and Greece). *Jasminum sambac* is also widely distributed in Asia but is mainly recorded in India. There are only five records of *J. sambac* in North-East Europe. We did not include *Trachelospermum* sp. as host plant, as the species has not been identified, and *Volkameria inermis* as it is a tropical species, essentially distributed on the seeshore in South East Asia between India and the South Pacific islands.

### Environmental variables

We used the 19 climatic variables available in WorldClim version 2.0 (www.worldclim.org) at a spatial resolution of 2.5 minutes. We also used the potential distribution of the four *Jasminum* species as biotic variable for *C. ayyari*. The procedure is explained below.

### Niche modelling

Before modelling the distribution of species, we analyse the correlation between variable using ‘Hmisc’ package and Pearson coefficient, either for the current and the future conditions. The sets of current and future climatic variables with a correlation coefficient up to 0.7 were different. As collinearity affects models trained on data at different times, we opted for variables issued from a principal component analysis (PCA) [17]. We did a PCA of the 19 climatic variables either of the current and future conditions using ‘FactoMineR’ package, and we selected the first nine PCA components as variables, accounting for 99% of the variance. *Jasminum* species modelled distributions were also used as biotic variables as they were not correlated up to 0.7 to the PCA components used as variables, and as *Jasminum* species and *C. ayyari* distributions were not correlated.

Data were cleaned before modelling using ‘CoordinateCleaner’ package [18], to get rid of inadequate occurrences such as the specimens deposited in museums, occurrences wrongly geo-referenced or duplicated. We considered the tests ‘urban_ref’, ‘capitals_ref’, as null since *Jasminum* species are ornamental plants and therefore can be present in urban environments. Pseudo-absences were added to the dataset of occurrences. We selected the same amount of pseudo-absences as occurrences as suggested by Bardet-Massin et al. [11]. We randomly selected 10 sets of pseudo-absences for each *Jasminum* species. Assuming that the distribution of *C. ayyari* is not in equilibrium [19] as we deal with an invasive species, we selected pseudo-absence out of a convex hull calculated on the basis of the known occurrences [20].

In a first step, we modelized the potential distribution of *Jasminum* species considering PCA variables, and projected it into the future (2081-2100 period). In a second step, we did a first modelling of *C. ayyari* potential distribution, considering PCA variables only, and we did a second one considering PCA variables and including the rasters resulting of the modelled distributions of *Jasminum* species as biotic variables. Prior to this last analysis, we tested the correlation between *C. ayyari* distribution and the one of *Jasminum* species. The two modelling were compared to see the influence of *Jasminum* species distribution on *C. ayyari* modelling. We projected the distribution of *C. ayyari* into the 2081-2100 period as well, using only PCA variables corresponding to the period, and in a second step including also the modelled projection into the future of *Jasminum* species. For the projection into the future, we considered SSP585 scenario, the worst scenario of global warming, corresponding to an additional radiative forcing of 8.5 W/m², and we use the CNRM-CM6-1 model, which presents particularly fine parameter calibration strategies tailored to the geographic area considered [21]. Modelling was made using ‘Biomod2’ package [22] and the following options:

All the models were tried except Maxent which is the only one based on presence only, i.e. Generalized Linear Model (GLM), Generalized Additive Model (GAM), Generalized Boosting Model (GBM), Classification Tree Analysis (CTA), Artificial Neural Network (ANN), Surface Range Envelop (SRE), Flexible Discriminant Analysis (FDA), Random Forest (RF), and Multiple Adaptive Regression Splines (MARS).

We run 10 evaluations, split the data in 20% test data and 80% modelling data. Models were selected to be combined using a threshold of 0.7 on the basis of TSS evaluation. Modelling was evaluated by combining all model, as suggested by [23], with an arbitrary threshold up to 0.7 and using the TSS binary metric as recommended by [24]. Combined model was evaluated using Boyce index with ‘ecospat’ package [25, 26].

### Niche comparison

We also compared ecological niches of *Jasmium* species and *C. ayyari* in the invaded area. Ecological niche estimates were performed with the ‘ecospat’ package [26, 27]. A Mantel test was done first to test spatial autocorrelation between variables. Niche overlap between native and invaded areas was also measured using the Schoener’s D index [27–29]. It can range from 0, for niches with no overlap, to 1, for completely identical niches. We evaluated niche overlap on the basis of the 19 climatic variables. We compare niche overlap only for *C. ayyari*, *J. grandiflorum* and *J. officinale*, as the two other *Jasminum* species do not have enough occurrences in the invaded area to be analysed.

We finally compared invasive vs. native niches of *C. ayyari* with and without *Jasminum* species modelling as biotic variables to see the influence of the host plants on the invasive potential of *C. ayyari*.

## Results

*Jasminum officinale* totalizes 1381 occurrences restricted to the areas studied, of which 78% are located in Europe. It totalizes 1065 occurrences after cleaning. *Jasminum sambac* totalizes 626 occurrences after cleaning over 724 existing. *Jasminum grandiflorum* totalizes 186 occurrences after cleaning over 203 existing, and *J.multiflorum* totalizes 27 occurrences after cleaining over 40 existing. *Corythauma ayyari* totalizes 48 occurrences, of which 37 are located in Europe and 11 are located in Asia. It totalizes 34 occurrences after cleaning (S1 Data) (Fig 1).

**Fig 1.**
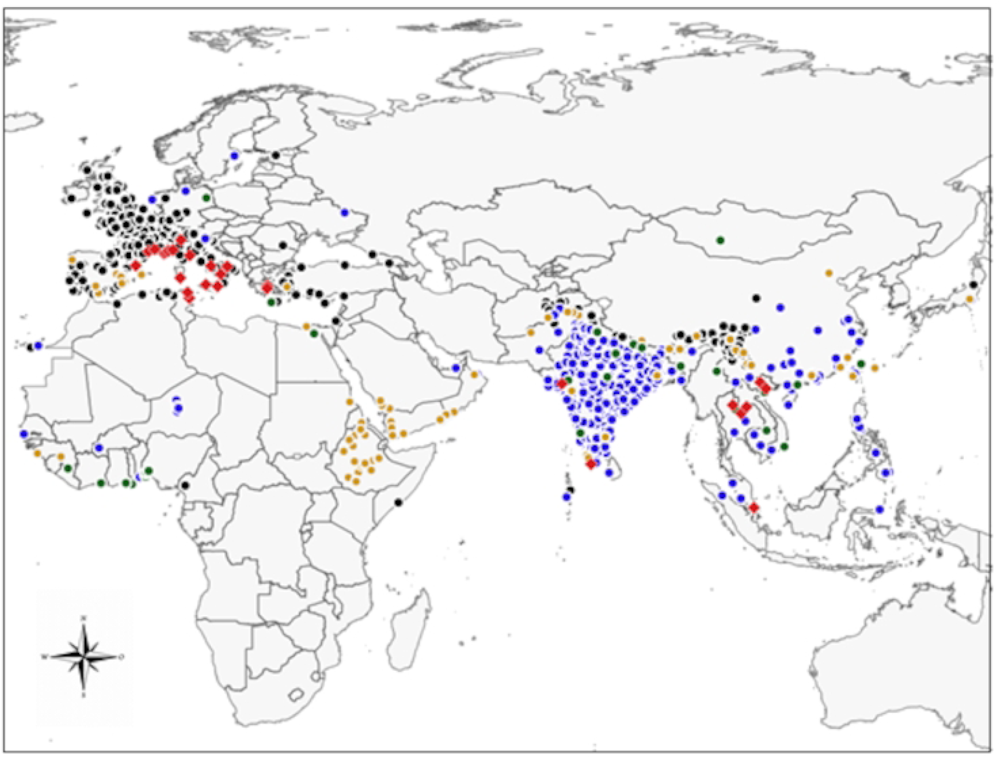
Map of the occurrences used. Map of the study area showing the occurrences of *C. ayyari* (red diamonds), and its four main host plants, *J. officinale* (black dots), *J. grandiflorum* (orange dots), *J. multiflorum* (green dots), and *J. sambac* (blue dots).

### Niche modelling

Boyce index is high for *J. officinale* (0.952), *J. grandiflorum* (0.946) but low for *J. multiflorum* (0.65), and *J. sambac* (0.668). Including *Jasminum* distributions as variable, Boyce index for *C. ayyari* distribution modelling is 0.760; while without *Jasminum* variables it is 0.659.

*Jasminum officinale* current distribution modelling shows a high suitability of habitat in all South West Europe, including Belgium, France, The Netherlands, Great Britain, Portugal, North and West sides of Spain, North and East Mediterranean coasts, including Maghreb coasts, and South coast of Black Sea (S1A Fig). High suitability areas include a thin band along the Himalayan range and part of Sichuan and Yunnan.

*Jasminum officinale* future projection shows a loss of suitable areas in most of eastern Europe, central regions of Spain, North of Great Britain and Irland, leaving a thin range along Mediterranean and Black Sea coasts (S1B Fig). It also shows a loss of the thin band along Himalayan range and the spot in Sichuan and Yunnan regions. It gains suitable areas in South East of China.

*Jasminum grandiflorum* current distribution modelling shows a high suitability all around the Mediterranean borders, extending to Portugal and the Spanish Atlantic coasts. In Asia, it includes the West side of India, central regions of India, a band along the Himalayan range, extending eastern to a region including Sichuan, Yunnan, Myanmar, and also East China with a lesser suitability (S1C Fig).

Projection into the future of *J. grandiflorum* distribution narrows the circum-Mediterranean band of suitable areas, extends to South and lowers the suitability in the Asian area (S1D Fig). Area of high suitability for *Jasminum multiflorum* is concentrated in South East Asia, and includes also West part of India and a band along the Himalayan range. Moderately suitable areas in North and West Europe include Norway, Ireland, Scotland, Wales, Bretagne in West of France, North West of the Iberic Peninsula, and a thin band along East coast of Adriatic Sea (S1E Fig).

Projection into the future of *J. multiflorum* distribution shows a restriction of the current distribution modelling, except the very North of Europe including Norway, Scotland, Ireland and Wales (S1F Fig).

*Jasminum sambac* current distribution modelling shows a suitable area restricted to India, the Himalayan range, South East China, Vietnam, Myanmar and some other areas into the South East Asia, comprising Thailand, Laos and Malaysia (S1G Fig).

Projection into the future of *J. sambac* distribution lowers the suitability of most of the area, and restricts it to a West region of India and Pakistan, and parts of Thailand and Myanmar (S1H Fig). Considering only PCA components, *Corythauma ayyari* current distribution modelling shows a high suitable habitat on the North Mediterranean coasts in Spain, France, Italy, the Croatian coasts, the Aegean and Marmara coasts, the South of Black Sea coast, part of Bretagne in France and at a lower degree parts of North and Central Europe except North Scandinavia. In Asia, suitable areas include India except West coast, Thailand, Myanmar, Laos, Vietnam and Cambodgia except West coast of South East Asia, Malaysia included, a large part of South and East China (Fig 2A).

**Fig 2.**
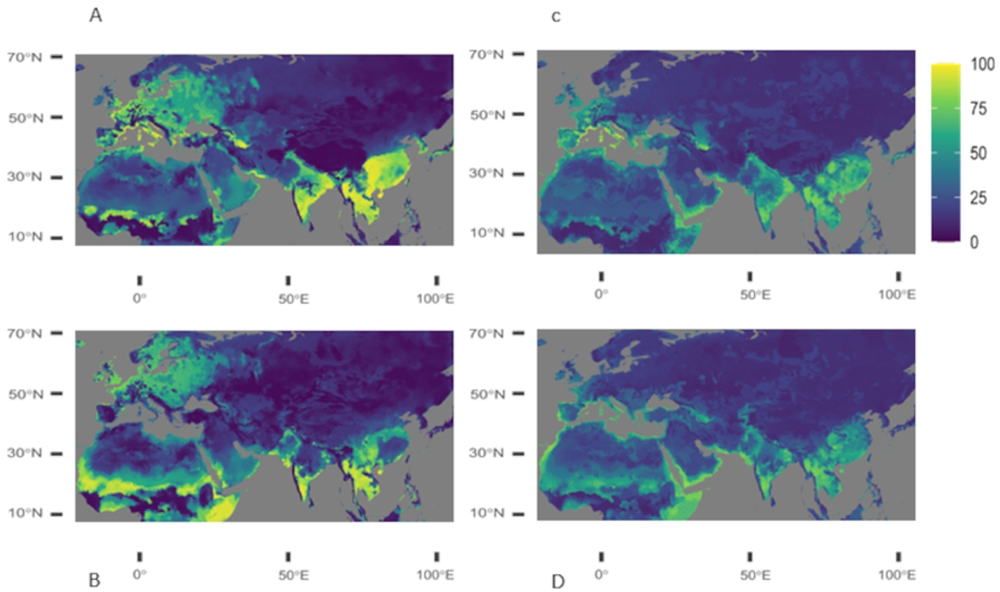
Projections of the current modeling and future projection of C. ayyari distribution. Projections of the current (upper row) and future (lower row) environmental conditions for *C. ayyari* using climatic variables only (left colon), and including also *Jasminum* species distribution as variables (right colon). The scale represents percentage of suitability along a gradient of colors.

*Corythauma ayyari* distribution projected into the future shows a reduction of the current suitable areas, also moving North of Europe facing the Channel and the North Sea where unsuitable areas turned 50% suitable (Fig 2B). In Asia, North India and East China turned unsuitable.

Correlation between *Jasminum* species distribution and *C. ayyari* distribution are low (ranging between 0.31 to 0.46, Table 1). Correlation between *Jasminum* species does not exceed 0.53 (between *J. multiflorum* and *J. grandiflorum*), lesser than the usual threshold accepted for correlated variables in niche modelling which is 0.7. As such, including *Jasminum* species distribution as variables does not bias modelling.

**Table 1.**
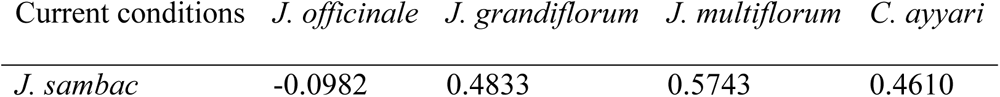
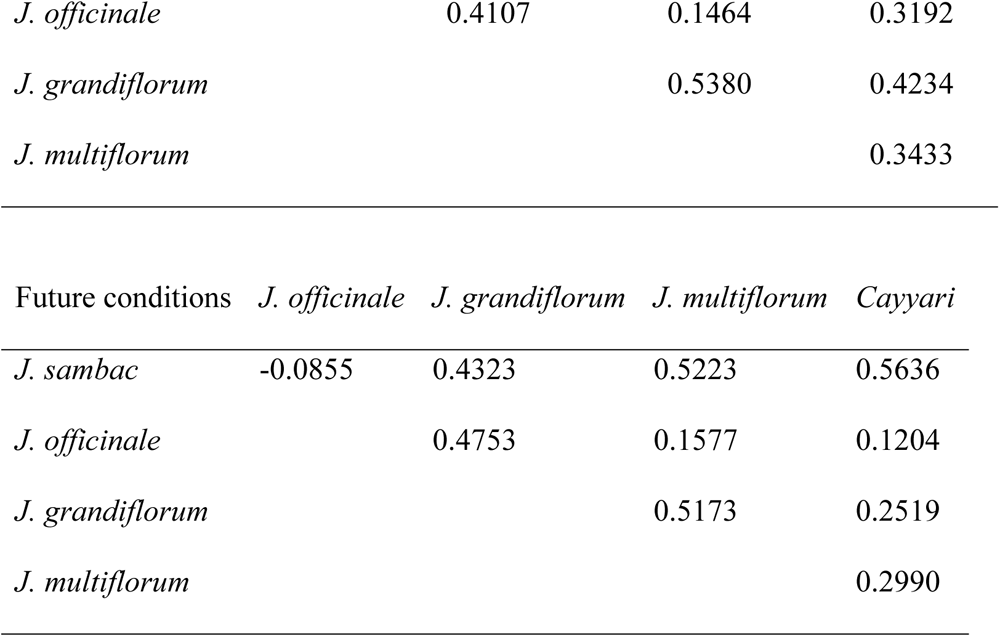
Correlations between species distribution.

Table 1 legend : Correlation between species distribution using Pearson coefficient, under current condition (above) and future conditions (below). Species distributions are rasters resulting from species distribution modelling using PCA components as variables.

When including *Jasminum* species distribution as variables, *C. ayyari* current distribution modelling shows suitable areas reduced to North Mediterranean coasts, including Aegean and Marmara coasts. It includes also part of West Europe and Maghreb coasts with a lesser suitability. In Asia, areas are not highly suitable, however, they include Part of South and East India, central South East Asia, except Malaysia and a large region in South East China (Fig 2C). *Corythauma ayyari* distribution projected into the future with Jasminum species distributions reduces suitable areas to a thin range along Mediterranean coasts, enlarging suitable areas to most of South Mediterranean coasts, Portugal, the Atlantic side of Spain and South West of France. Suitability of areas is reduced in Asia, except in West India in Gujarat region and Pakistan (Fig 2D).

The most contributing variables for *C. ayyari* modelling, all models together are the 5^th^, 1^st^ and 2^nd^ PCA components (39%, 15% and 12%, respectively). When including *Jasminum* distribution as variables, the three most contributing variables are the 5^th^ PCA component (28%) and *J. officinale* and *J. grandiflorum* distribution modeling (23% and 13%, respectively) (S1 table).

### Niche comparison

Overlap of *Jasminum* species and *C. ayyari* niches in the native area is low between all species, except between *J. multiflorum* and *J. grandiflorum* (70%), for which the similarity is significant (Table 2).

**Table 2.** Niche overlap, equivalency and similarity tests between species in the native area.

| Overlap | *J. grandiflorum* | *J. multiflorum* | *J. sambac* | *C. ayyari* |
| --- | --- | --- | --- | --- |
| *J. officinale* | 0.0510 | 0.0385 | 0.0837 | 0.0707 |
| *J. grandiflorum* |  | 0.7010 | 0.1608 | 0.0156 |
| *J. multiflorum* |  |  | 0.2427 | 0.0329 |
| *J. sambac* |  |  |  | 0.1138 |
| Similarity | *J. grandiflorum* | *J. multiflorum* | *J. sambac* | *C. ayyari* |
| *J. officinale* | 0.3168 | 0.4158 | 0.1881 | 0.1287 |
| *J. grandiflorum* |  | 0.0297 | 0.1188 | 0.3762 |
| *J. multiflorum* |  |  | 0.0792 | 0.3168 |
| *J. sambac* |  |  |  | 0.1089 |

Table 2 legend. Niche overlap, similarity and equivalency test between species in the native area.
| Equivalency | *J. grandiflorum* | *J. multiflorum* | *J. sambac* | *C. ayyari* |
| --- | --- | --- | --- | --- |
| *J. officinale* | 1 | 1 | 1 | 1 |
| *J. grandiflorum* |  | 0.3636 | 1 | 1 |
| *J. multiflorum* |  |  | 0.3636 | 0.6363 |
| *J. sambac* |  |  |  | 0.5454 |

Table 2 legend. Niche overlap, similarity and equivalency test between species in the native area. Significant values are in italic.

Overlap between species in the invaded niche is significantly similar only between *J. officinale* and *C. ayyari*. It has not been tested for *J. multiflorum* and *J. sambac* due to the too low number of occurrences in Europe (Table 3).

**Table 3.**
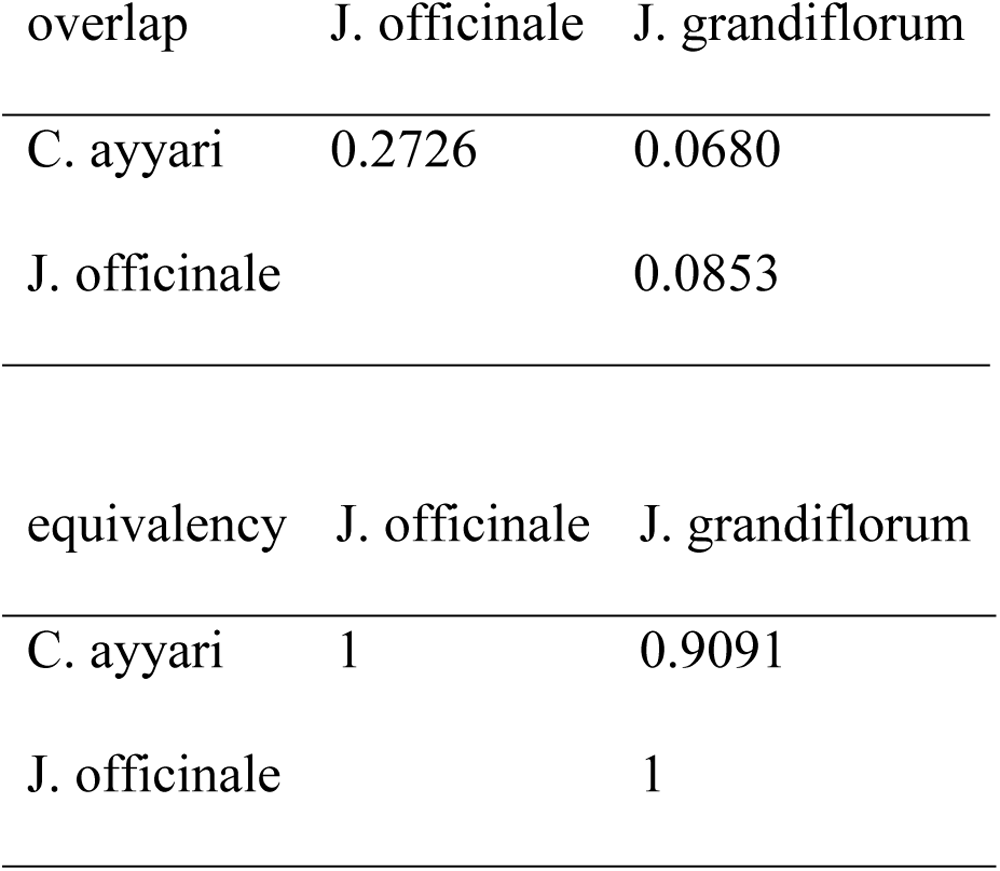
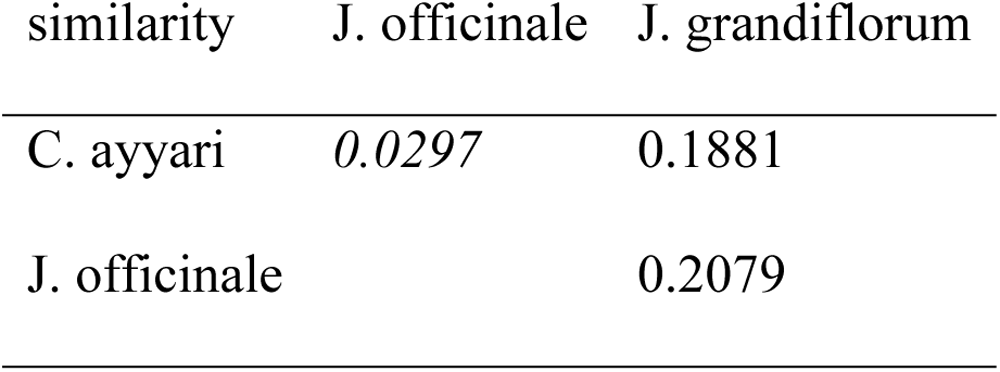
Niche overlap, equivalency and similarity tests between species in the invaded area.

Table 3 legend. Niche overlap, similarity and equivalency test between species in the invaded area. Significant values are in italic.

Overlap between European and Asian niches of *J. officinale* is high (Schoener’s D=0.1670). Niches are equivalent but not similar (P value= 0.0198 and 0.1188, respectively). Overlap between native and invaded niches of *J. grandiflorum* is low (Schoener’s D = 0.0138), however it is similar (P value=0.0396) but not equivalent (P value =1). Comparison of native and invaded niche for *C. ayyari* shows a very low overlap (D=0.024). It is neither equivalent, nor similar (P value=0.9604 and 0.0594, respectively).

## Discussion

*Corythauma ayyari*, seems to thrive in the warm-summer Mediterranean climate (Csa) of the European regions [30–32]. This temperate climate (minimum temperatures between −3°C and 18°C throughout the year) offers dry and warm summers, with precipitation less than 40 mm and maximum temperatures above 22°C, and mild winters [31]. The period of greatest activity of *C. ayyari*, would probably occur between late spring and early autumn, coinciding with the warm summers of the Mediterranean climate, while in winter during the coldest periods, *C. ayyari* would overwinter [7, 32, 33].

The native distribution of *C. ayyari* is restricted to tropical climates (minimum temperature greater than or equal to 18 °C) of equatorial savannah with dry winter (rainfall in the driest month less than 60 mm) (Aw), and to warm temperate climates with dry winter and hot summer (average temperature of the hottest month greater than 22 °C) (Cwa) [31, 32]. The temperature and the presence of a dry period are thus important variables common to both native and invaded areas of *C. ayyari*. Low temperatures seem to be the main constraints to the establishment of *C. ayyari*. Thermophilic in nature, it survives, like most other Tingidae, at a temperature range of 15°C to about 40°C, with an optimal development and fecundity rate around 30°C [4, 5, 33–35]. At lower temperatures, it enters hibernation. The mild winter of the Mediterranean climate should therefore be favourable to it. Thus, under the worst-case scenario ssp585, temperatures will increase by 3.3°C to 5.7°C, for the period 2081-2100 [36]. This increase in temperature in the Mediterranean climate may facilitate optimal development of *C. ayyari*, while reducing its hibernation periods in future years.

*Jasminum* species are mainly recorded from their native area in Asia, except *J. officinale* which is mainly recorded from Europe, due to its importation as ornamental plant. Then, distribution modelling shows that Europe is mainly suitable for *J. officinale*, but weakly suitable for *J. multiflorum*, and not suitable for *J. sambac*. Mediterranean coasts only are suitable for *J. grandiflorum*.

Not surprisingly, the main suitable areas for *C. ayyari* are in the native region, South East Asia, despite few occurrences recorded there, comparing to the occurrences in Europe. Outside its original range, according to our models when considering climatic variables only, the most suitable areas for *C. ayyari* are concentrated on the North coasts of Mediterranean Sea, where it has been recently recorded, and where *J. officinale* and *J. grandiflorum* are present. Parts of North West of Europe remain also suitable for *C. ayyari*. Not surprisingly also, suitable areas are restricted to a thinner range along North Mediterranean coasts and part of Western Europe at a lower suitability, when *Jasminum* species are included as variables. This is mainly due to *J. officinale* and *J. grandiflorum* distributions, as *J. sambac* has no suitable areas in Europe, and *J. multiflorum* has a low suitability in Northern Europe. As expected, *C. ayyari* suitable area is restricted by its host plants distribution.

Projections into the future for *J. grandiflorum and J. officinale* show that suitable areas will be restricted to a narrower band along Mediterranean coasts and will also move West of Europe. In consequence, *C. ayyari* will also be restricted to the same areas, according to the projection including *Jasminum* species as variable. Otherwise, *C. ayyari* suitable areas will extend North and East of Europe.

Modelling of *C. ayyari* distribution and projection into the future differs when including or not *Jasminum* species as variable. *Jasminum* distribution modelling shows that *C. ayyari* will not benefit from its main host plants to spread out in Europe as would suggest *C. ayyari* modelling using only climatic variable. In addition, including the host plants as biotic variables render the modelling more accurate (see Boyce index). This show the importance of integrate biotic variables such as host plants for phytophagous species in modelling.

At the present day, the suitable area of *C. ayyari* in Europe does not fit with *J. officinale* distribution and is restricted to Mediterranean coasts, like *J. grandiflorum*. Harbours are the main entrance of the host plants, and then *C. ayyari* would have been introduced in Europe with its host plants [4]. It was recorded on *J. grandiflorum* in Spain and Syria; *J. sambac* and *J. grandiflorum* in Tunisia; *J. sambac* and *J. multiflorum* in Israel; *J. officinale* and *J. grandiflorum* in Italia. However, the invaded niche of *C. ayyari* and of *J. officinale* are similar (P value=0.0297), while this is not the case of *J. grandiflorum*’s niche, and Europe is not suitable for *J. sambac*. As native and invaded niche of *C. ayyari* is neither equivalent nor similar, this would suggest that *C. ayyari* switched to one niche to another, and could potentially benefit from the presence of *J. officinale* to setting up in Europe. More South in Mediterranea, *J. sambac* and *J. grandiflorum* could stay the main plant hosting *C. ayyari*.

Niche shift of introduced plant species is not rare as stated by Atwater et al. [37]. This assumption highly depends on variable use and modelling method and has been debated on invasive plants [38, 39], and also on introduced mammals [13].

The adaptability of *C. ayyari* suggest that it could spread to other host plants. Indeed, *C. ayyari* has been observed on the genera *Lantana*, *Ocimum*, *Musa*, *Hedychium*, and *Trachelospermum*, and the species *Althea officinalis*, *Daedalacanthus nervosus*, and *Volkameria inermis* (Novoselsky & Freidberg 2013; Carapezza. 2014; Pedata et al. 2014, Haouas et al. 2015). The observation of *C. ayyari* on these plants does not mean they are host plants but may be only visited plants. Most of them are popular ornamental plants and widespread in gardens so, their trade would be the main source of the introduction of *C. ayyari* and one of the major factors for its expansion. For example, *Trachelospermum jaminoides* is probably the *Trachelospermum* species on which *C. ayyari* was found in France (Streito et al. 2010), as it is the only species of the genus recorded in Europe. It has a wide distribution in Europe and could be a potential host, facilitating *C. ayyari* invasion.

## Conclusions

This study shows that host plants distribution is key to phytophagous species. They may restrict invasive potential of the invader. However, oligo-and polyphagous species have the possibility to switch to several host plants and then invade new areas, switching from one niche to another. *Halyomorpha halys* (Stal, 1855) is a typical example of a successful invasive species as it is highly polyphagous and feeds on more than 120 wild or cultivated plant species (Streito et al. 2021). Modelling distribution depends on data availability, as invasive species provide scarce data when early considered. Therefore, modelling their potential invasiveness is often done once species already established and disrupted native ecosystem.

In several countries like India, Pakistan, Egypt, Tunisia, *Jasminum*, especially *J. grandiflorum* and *J. sambac*, have a very high symbolic value and are emblematic plants of the country (Haouas et al. 2015). They are also used in the perfume industry. The invasion of *C. ayyari* in these regions would cause significant economic but also symbolic damage. Therefore, as human activity is the main source of *C. ayyari* expansion due to the ubiquitous presence of ornamental plants in gardens and parks, means of management in the latter are conceivable, and would be different from a widely cultivated plant as food resource.

## Acknowledgements

The authors thanks Alexandre Schickele for providing help with R scripts, and Jean-Claude Streito for fruitful comments on the manuscript.

## Supporting information

**S1 Data. GPS coordinates of occurrences of the species used in the analyses**. GPS coordinates of *Jasminum officinale* (Joff), *J. grandiflorum* (Jgra), *J. multiflorum* (Jmul), *J. sambac* (Jsam) and *C. ayyari* (Cay) occurrences after cleaning (see method section), used for analyses.

**S1 Fig. Projections of the current (on the left) and future (on the right) environmental conditions resulting from the niche modelling for *Jasminum* species**. Ensemble projection combines all model with an arbitrary threshold up to 0.7 and uses the TSS binary metric, for the four main *Jasminum* species hosting *C. ayyari* : A, B) *J. officinale*; C, D) *J. grandiflorum*; E, F) *J. multiflorum*; G, H) *J. sambac*.

**S1 Table. Contribution of variables used for modelling *C. ayyari* distribution**. A first modelling uses only the first nine climatic variables issued from a PCA ; a second modelling includes also the four *Jasminum* species distribution as variable.

## Notes

### Competing Interest Statement

The authors have declared no competing interest.

